# Tropomyosin 1 deficiency facilitates cell state transitions to enhance hemogenic endothelial cell specification during hematopoiesis

**DOI:** 10.1101/2023.09.01.555861

**Authors:** Madison B Wilken, Gennadiy Fonar, Catriana Nations, Giulia Pavani, Victor Tsao, James Garifallou, Joanna Tober, Laura Bennett, Jean Ann Maguire, Alyssa Gagne, Nkemdilim Okoli, Paul Gadue, Stella T Chou, Nancy A Speck, Deborah L French, Christopher S Thom

**Author notes:** Correspondence: Christopher S Thom, Children’s Hospital of Philadelphia, 10-052 Colket Translational Research Building, 3501 Civic Center Blvd, Philadelphia, PA 19104. Contributed equally (Madison B Wilken, Gennadiy Fonar).

## Abstract

Tropomyosins coat actin filaments and impact actin-related signaling and cell morphogenesis. Genome-wide association studies have linked *Tropomyosin 1* (*TPM1*) with human blood trait variation. Prior work suggested that *TPM1* regulated blood cell formation in vitro, but it was unclear how or when *TPM1* affected hematopoiesis. Using gene-edited induced pluripotent stem cell (iPSC) model systems, *TPM1* knockout was found to augment developmental cell state transitions, as well as TNFα and GTPase signaling pathways, to promote hemogenic endothelial (HE) cell specification and hematopoietic progenitor cell (HPC) production. Single-cell analyses showed decreased *TPM1* expression during human HE specification, suggesting that *TPM1* regulated in vivo hematopoiesis via similar mechanisms. Indeed, analyses of a *TPM1* gene trap mouse model showed that *TPM1* deficiency enhanced the formation of HE during embryogenesis. These findings illuminate novel effects of *TPM1* on developmental hematopoiesis.

## Introduction

*Tropomyosin 1* (*TPM1*) is one of four mammalian *Tropomyosin* genes (*TPM1-4*) that bind virtually all cellular actin to regulate cell shape, strength, and molecular signaling ^1,2^. *TPM* genes produce >40 protein isoforms, each of which can differentially impact actin filament structure and cellular dynamics ^2,3^. For example, some high molecular weight *TPM1* isoforms (e.g., 1.6/1.7) associate with actin stress fibers that typically promote cell adhesion ^2^, whereas low molecular weight *TPM1* isoforms (e.g., 1.8/1.9) promote lamellipodial persistence and cell motility ^4^. TPM1 activities are known to impact neuronal, cardiac, and ocular tissue development ^5–7^. Genome-wide association studies (GWAS) have implicated *TPM1*-associated polymorphisms with human blood trait variation ^8–10^, suggesting that *TPM1* may also regulate blood cell formation and/or function.

Hematopoiesis is a highly orchestrated process by which embryonic endothelial cells develop into specialized ‘hemogenic’ endothelial cells (HE), which subsequently produce the hematopoietic stem and progenitor cells (HPCs) that support mature blood cell formation throughout the mammalian lifespan ^11^. The first HPCs are produced in the embryonic yolk sac during primitive hematopoiesis. Later, HPCs are produced in a number of locations including the dorsal aorta-gonad-mesonephros region during definitive hematopoiesis. Although the stages of hematopoietic development are well characterized, the inability to efficiently recapitulate blood formation in vitro demonstrates that some factors remain unknown.

Endothelial specification into arterial, venous, and HE subtypes occurs at the onset of cardiac function and pulsatile blood flow in the embryo ^12^. Factors that regulate HE specification are not fully understood, but recent data have shown that this process requires coordinated retinoic acid, cKit, and Notch pathway signaling, as well as tight cell cycle control ^13–15^. Meis1 activity also helps to establish HE identity ^16^, as do proinflammatory signals from TNFα that activate Notch and NFκB signaling pathways to establish the HPC fate ^17^. HE are marked by expression of RUNX1, which cooperates with TGFβ signaling to regulate HPC formation ^18^. During both primitive and definitive hematopoiesis, coordinated transcriptional, signaling, and structural changes prepare HE cells to undergo a dramatic morphogenesis from planar, adherent cell types into spherical, non-adherent HPCs. This cell state change from HE to HPC is termed the endothelial-to-hematopoietic transition (EHT) ^19^. HE specification and EHT also occur in cell culture systems that model hematopoiesis.

HE specification and EHT resemble mesenchymal-to-epithelial and epithelial-to-mesenchymal cell state transitions (EMTs), respectively ^18^. Tropomyosins are known to regulate cell state transitions, including EMTs, in a number of tissue and physiologic states ^20^. Increased *TPM1* expression has been observed in cells undergoing EMT in the murine eye lens epithelium ^7^, and concurrent *TPM1* and *TPM2* deletion inhibited normal eye lens formation ^6^. *TPM1* deficiency has also been linked with EMT during cancer progression, as well as increased proliferation and migration in cell lines designed to solid tumor models ^21–25^. For example, *TPM1* mediates TGFβ-induced migratory behavior via actin cytoskeletal rearrangements and stress fiber formation in cultured epithelial cells ^23^. *TPM1* also constrains TNFα-mediated inflammatory signaling to regulate cultured arterial endothelial actin organization, migration, and proliferation ^26^. Actin cytoskeletal dynamics ^27^, TGFβ ^18^, and TNFα signaling ^17^ also regulate hematopoiesis, suggesting that *TPM1* might regulate blood formation through similar mechanisms.

We previously showed that *TPM1* normally constrains HPC formation in vitro from cultured induced pluripotent stem cells (iPSCs) ^10^, but it was unclear how or when *TPM1* impacted hematopoietic development. We reasoned that defining the mechanisms and developmental stages by which *TPM1* regulated hematopoiesis would provide novel approaches or complement existing strategies to enhance in vitro blood cell formation, and contribute to a broader understanding of the role that tropomyosins play in regulating cell development. We hypothesized that *TPM1* may regulate HE specification and/or EHT during hematopoiesis, given established links between *TPM1* and cell state transitions in other developmental systems. We found that *TPM1* was normally downregulated during hematopoiesis, and that constitutive *TPM1* deficiency promoted HE formation, with concomitant changes in inflammatory and GTPase signaling, without compromising HPC function. Using single-cell transcriptomics, we also found that *TPM1* expression was diminished in HE compared to stromal and epithelial cells. Murine studies confirmed that *TPM1* haploinsufficiency increased HE specification in vivo, demonstrating that inhibition of this cytoskeletal regulatory molecule is sufficient to promote HE formation. These findings define a novel role for *TPM1* in hematopoiesis across mammalian species and developmental ontogeny.

## Results

### Tropomyosin 1 deficiency enhances in vitro endothelial cell formation without augmenting cell cycle kinetics

An induced pluripotent stem cell (iPSC) model system was used to study the role of *TPM1* during in vitro primitive hematopoiesis, which includes defined iPSC, mesoderm, endothelial, and HPC stages of development ^10^ (**Fig. 1A**). Our first goal was to define when TPM1 protein was expressed in this system. We confirmed that high molecular weight TPM1 protein (e.g., TPM1.6/1.7) was expressed in adherent cell types, including iPSC-derived mesoderm and endothelial cells, but was virtually abolished in non-adherent HPCs (**Fig. 1B**). The absence of high molecular weight TPM1 protein was contrasted by the persistence of low molecular weight TPM1 isoforms (e.g., TPM1.8/TPM1.9) in HPCs and mature blood cell types (**Supplementary Fig. 1S1**). The dynamic change in expression suggested that high molecular weight TPM1 might impact adherent cell biology or development, including endothelial cells.

**Figure 1.**
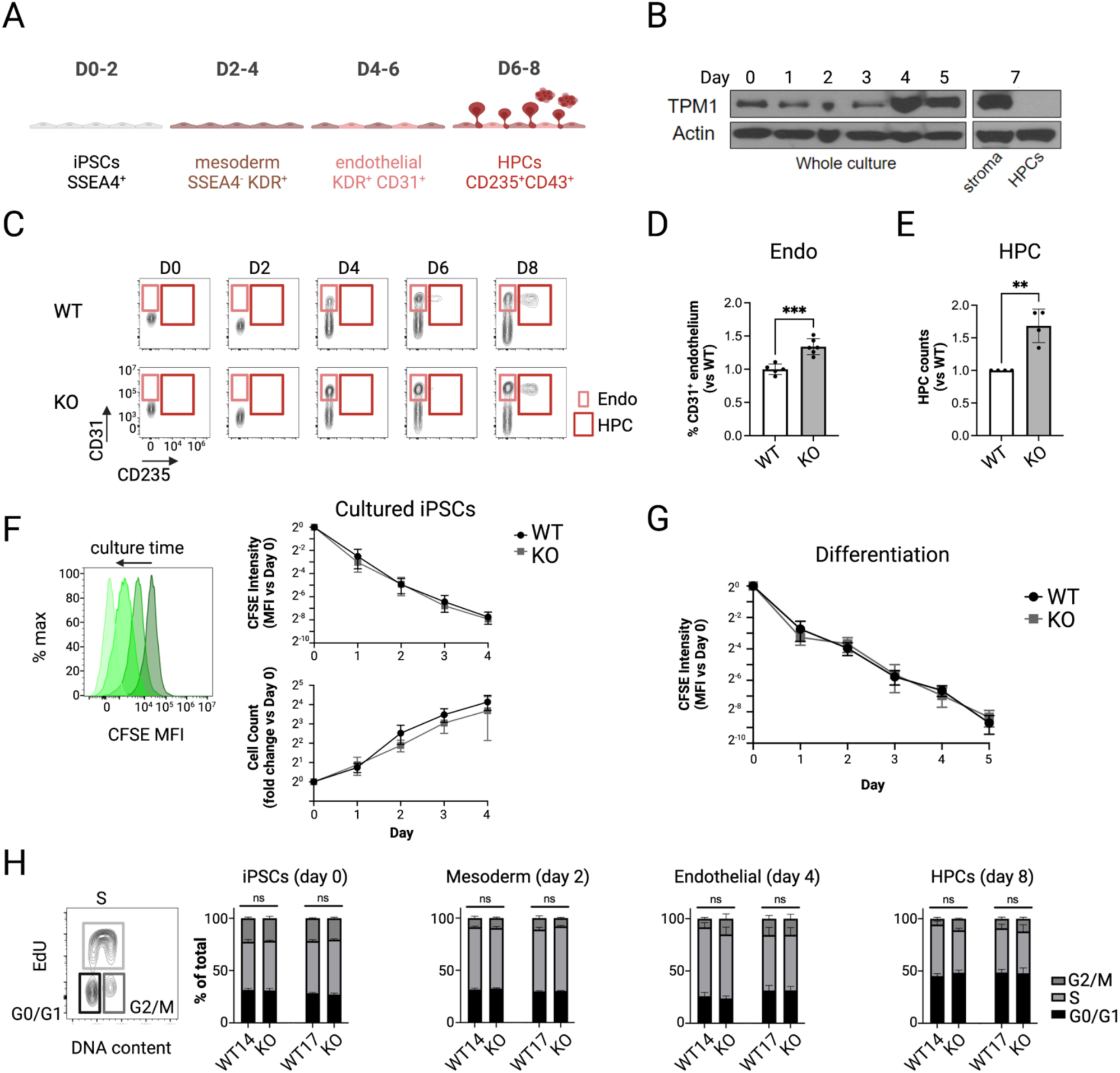
*Tropomyosin 1* deficiency enhances in vitro endothelial and hematopoietic progenitor cell (HPC) formation without enhancing proliferation. (A) Schematic overview of primitive in vitro hematopoiesis, including relevant cell types and cell surface markers over the differentiation timeline. (B) Western blot showing TPM1.6/1.7 protein expression over the course of in vitro hematopoiesis. On day 7, adherent stromal cells and non-adherent HPCs were collected and analyzed separately. (C) Exemplary flow cytometry plots during differentiation, with boxes highlighting CD31^+^ endothelial cells and CD235^+^ HPCs in wild type (WT) and *TPM1KO* (KO) cultures. (D) *TPM1KO* cultures show enhanced CD31^+^ endothelial cell percentage (%) vs isogenic WT cultures on days 4-6 (CHOP17 cell lines). Bar plots show mean±SD, significance assessed by t test. (E) *TPM1KO* cultures produce more non-adherent CD235^+^ HPCs on day 7-8 than isogenic WT controls (CHOP17). Bar plots show mean±SD. (F) Mean fluorescence intensity (MFI) for CFSE, a non-toxic cell permeable dye, diminishes with each cell division in culture. CFSE MFI diminishes identically in cultured isogenic WT and *TPM1KO* iPSCs (CHOP10). Similarly, there were no significant differences in WT vs *TPM1KO* cell expansion over time by direct cell counting (n=3 experiments). (G) CFSE MFI diminishes identically in isogenic WT and *TPM1KO* cells undergoing primitive hematopoietic differentiation (CHOP10). (H) EdU-based analysis allows identification of cells in G0/G1, S, and G2/M cell cycle stages. Analysis of *TPM1KO* cells at the iPSC, day 2 SSEA4^-^ (mesoderm), day 4 CD31^+^ (endothelial), and day 8 CD235^+^ non-adherent HPC stages showed no significant differences in cell cycle progression compared with isogenic WT controls (n=4-9 per group). Bar plots show mean±SEM. *p<0.05, **p<0.01, ****p<0.0001 by ANOVA.

Next, we analyzed developmental progression in genome-edited *TPM1* knockout (KO) iPSCs that lacked all *TPM1* isoforms, compared with isogenic wild type controls^10,28^. *TPM1KO* cells showed normal developmental kinetics and cell surface marker expression, with increased endothelial cell and HPC yields compared to controls (**Fig. 1C-E** and **Supplementary Fig. 1S2**). These results confirmed our previous findings ^10^ and were consistent using three iPSC lines of different genetic backgrounds ^10,28^. We therefore used these 3 stem cell lines interchangeably throughout this study to investigate mechanisms of action.

We reasoned that *TPM1KO* could increase endothelial and HPC production by enhancing proliferation ^29^ or by enhancing HE specification ^19,20^ (**Fig. 1A**). To determine if *TPM1KO* altered cell proliferation, we examined cell permeable carboxyfluorescein succinimidyl ester (CFSE) washout (**Fig. 1F**). CFSE staining decreased identically in *TPM1KO* and isogenic WT controls, confirming that *TPM1KO* cells proliferated normally during differentiation (**Fig. 1F-G** and **Supplementary Fig. 1S3**). To determine if cell cycle progression was altered at specific developmental stages, we stained relevant cell populations with 5-Ethynyl-2’-deoxyuridine (EdU). *TPM1KO* cell cycle kinetics did not significantly differ from isogenic controls at any stage of development (**Fig. 1H** and **Supplemental Fig. 1S4**). These findings suggested that mechanisms other than increased cell proliferation were responsible for increased endothelial and hematopoietic cell production in cultured *TPM1KO* cells.

### Tropomyosin 1 deficiency increases expression of EMT and HE-related signaling pathways during in vitro hematopoiesis

We reasoned that *TPM1KO* might enhance specification of HE or other HPC precursors, or more generally facilitate cell state transitions (EMT/EHT), to promote hematopoiesis in the absence of increased proliferation. To ascertain if and when we could detect transcriptional evidence of developmental perturbations in cultured *TPM1KO* cells, we performed bulk RNA sequencing (RNAseq) analysis on defined cell types during in vitro hematopoiesis. We initially looked for evidence of global developmental perturbation in *TPM1KO* cells. Clustering analysis showed that *TPM1KO* gene expression generally matched isogenic wild type controls at each stage of differentiation (**Fig. 2A** and **Supplementary Fig. 2S1**). This occurred despite changes in actin- and focal adhesion-related gene expression in most *TPM1KO* cell types (**Fig. 2B, Supplementary Tables 1-4**). These findings supported the notion that *TPM1KO* cultures underwent normal developmental stage progression, albeit with expected differences in actin regulatory processes.

**Figure 2.**
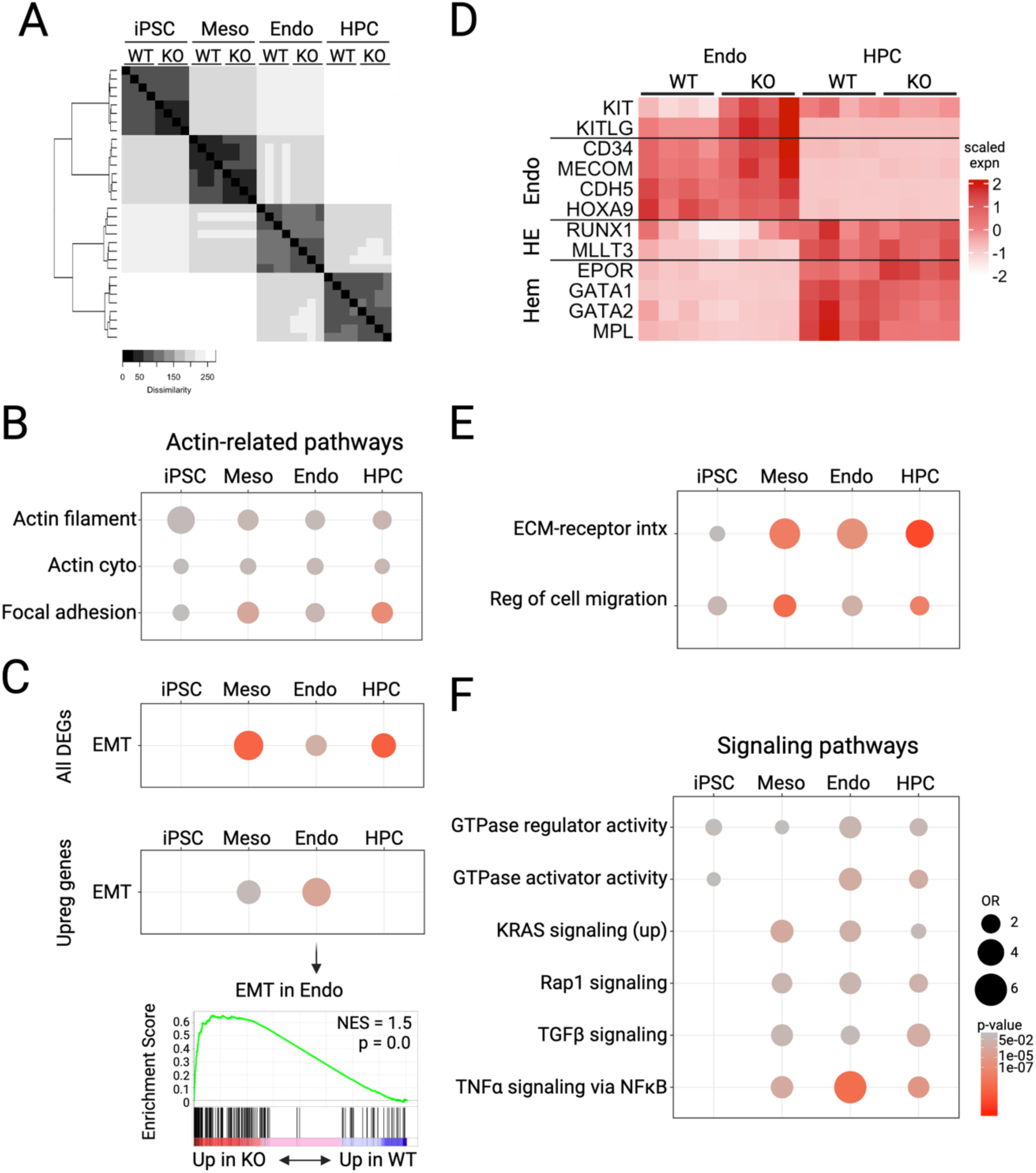
*TPM1* deficient cultures show normal maturation with enhanced epithelial-to-mesenchymal transition (EMT)-related gene expression. (A) Heap map demonstrates similarity of isogenic CHOP14 WT and *TPM1KO* samples at the iPSC, mesoderm, endothelial, and HPC stages of development. (B) Pathway analyses at the specified stages reveal changes in actin and focal adhesion pathways over the course of development. Bubble plots depict enrichment odds ratios (OR) and color reflects statistical enrichment (p-values) for the indicated pathways in *TPM1KO* vs isogenic WT control cells. (C) EMT-related gene expression is altered in *TPM1KO* cells at the mesoderm, endothelial, and HPC stages. EMT genes were specifically upregulated in *TPM1KO* endothelial cells. By Gene Set Enrichment Analysis, *TPM1KO* increases an EMT signature in endothelial cells. NES, normalized enrichment score. (D) Select endothelial, HE, and hematopoietic (Hem) gene expression in WT and *TPM1KO* cells at the endothelial and HPC stages. (E) Pathway analyses at the specified stages reveal changes in extracellular matrix-receptor interactions (ECM-receptor intx) and cell migration over the course of development. Bubble plots depict enrichment odds ratios (OR) and color reflects statistical enrichment (p-values) for the indicated pathways in *TPM1KO* vs isogenic WT control cells. (F) Pathway analyses at the specified stages reveal changes in signaling pathways over the course of development. Bubble plots depict enrichment odds ratios (OR) and color reflects statistical enrichment (p-values) for the indicated pathways in *TPM1KO* vs isogenic WT control cells.

We next assessed if and when EMT or related EHT gene expression changed in *TPM1KO* cultures. At the endothelial stage, we observed an increase in EMT-related gene expression by Gene Set Enrichment Analysis (**Fig. 2C**). This provided evidence that *TPM1KO* endothelium differed even at a relatively early time point, 2-3 days before HPC emergence. Changes in EMT gene expression could potentially indicate increased HE specification and preparation for EHT. The timing of collection (day 4) preceded full induction of RUNX1 and the downstream hematopoietic program, so we did not detect widespread changes in blood genes or pathways (**Fig. 2D**). However, *TPM1KO* endothelial cells did show increased *KIT* gene expression, which is important for HE specification ^15^ (**Fig. 2D**).

We envisioned two non-mutually exclusive mechanisms by which *TPM1KO* and actin cytoskeletal perturbations could enhance HE specification. First, *TPM1KO* could promote HE escape from the adherent endothelial cell environment to form HPCs ^19^ (**Fig. 1A**). Consistent with a biophysical mechanism, *TPM1KO* altered the expression of genes that impact extracellular matrix-receptor interactions and cell migration (**Fig. 2E** and **Supplementary Tables 1-4**). Second, altered actin dynamics could change the scaffolding necessary for signaling pathway regulation, including pathways necessary for HE specification and HPC formation ^30^. Consistent with signaling changes, we noted mild alterations in KRAS and Rap1 GTPase signaling, generally beginning at the mesoderm stage, as well as an overall increase in GTPase activation activity during *TPM1KO* development (**Fig. 2F** and **Supplementary Tables 1-4**). We also noted modest changes in TGFβ signaling, along with changes in TNFα signaling via NFκB that were most evident in *TPM1KO* endothelial cells (**Fig. 2F**). TGFβ ^18^, TNFα ^17^, and GTPase signaling mechanisms ^31^ can each promote HE specification and/or EHT. Taken together, these findings suggested that *TPM1KO* promoted hematopoiesis through multiple mechanisms, including altering actin dynamics, physical cell interactions, and signaling activities in developing endothelial cells.

### Tropomyosin 1 deficiency enhances HE specification to produce functional HPCs

We next sought functional evidence to determine if *TPM1KO* facilitated HE specification and/or EHT. To determine if *TPM1KO* increased the frequency of HE cells that undergo EHT, we cultured sorted day 4-5 CD31^+^CD43^-^ endothelial cells, plated them in limiting dilution in hematopoietic cytokines, and subsequently quantified CD43^+^ HPCs in each well ^18^ (**Fig. 3A**). These experiments identified an increased frequency of HE in *TPM1KO* cultures in two independent iPSC lines (**Fig. 3B**).

**Figure 3.**
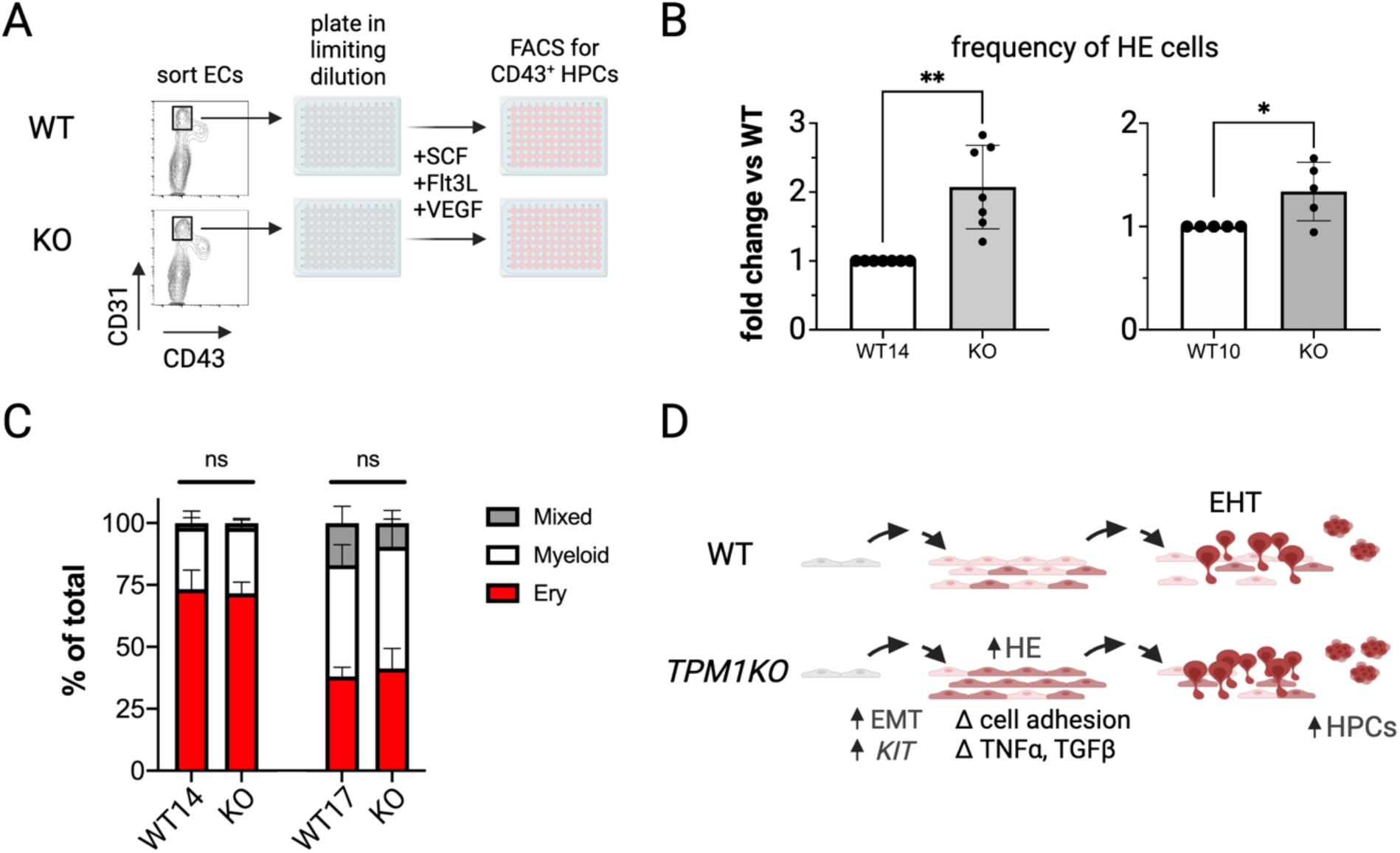
Tropomyosin 1 deficiency enhances HE cell specification during in vitro hematopoiesis. (A) To set up limiting dilution assays, CD31^+^CD43^-^ endothelial cells from *TPM1KO* and isogenic WT controls were sorted and plated in limiting dilution. After culturing in hematopoietic cytokines, the quantity of CD43^+^ HPCs was quantified by FACS. (B) The frequency of HE cells in *TPM1KO* vs isogenic WT cultures were quantified and normalized to WT frequency in each experiment (n=5-7 samples over 3-5 experiments). *p<0.05, **p<0.01. (C) In colony formation assays, *TPM1KO* and isogenic WT controls produce similar numbers of erythroid, myeloid, and mixed colonies (n=6-8 assays per group). (D) During primitive in vitro hematopoiesis, *TPM1KO* alters EMT, signaling pathways, and gene expression to enhance yields of HE (brown) and functional HPCs.

Although *TPM1KO* cultures increased the frequency of HE cells and HPCs, we wanted to confirm whether *TPM1KO* HPCs had any functional limitations or lineage bias. In colony formation assays, we found normal quantitative and qualitative production of erythroid, myeloid, and megakaryocyte colonies (**Fig. 3C** and **Supplementary Fig. 3S1**). These results complemented prior findings that showed normal function in *TPM1KO* megakaryocytes cells ^10^. Hence, *TPM1KO* enhanced in vitro hematopoiesis by increasing HE specification and functional HPC yields (**Fig. 3D**).

### Tropomyosin 1 expression changes during in vivo HE specification and EMT/EHT

To determine whether *TPM1* also regulated in vivo hematopoietic development, we analyzed recently published single cell RNA sequencing (scRNA) data sets that have provided insights into the human and mouse hematopoietic systems ^32,33^. Among profiled human embryonic cells, we identified high *TPM1* expression in both stroma and epithelial subsets and lower *TPM1* expression in other populations, including HE (**Fig. 4A**). Reduced *TPM1* expression in HPCs matched our in vitro data (**Fig. 1B**). This finding suggested that *TPM1* is normally diminished during HE specification.

**Figure 4.**
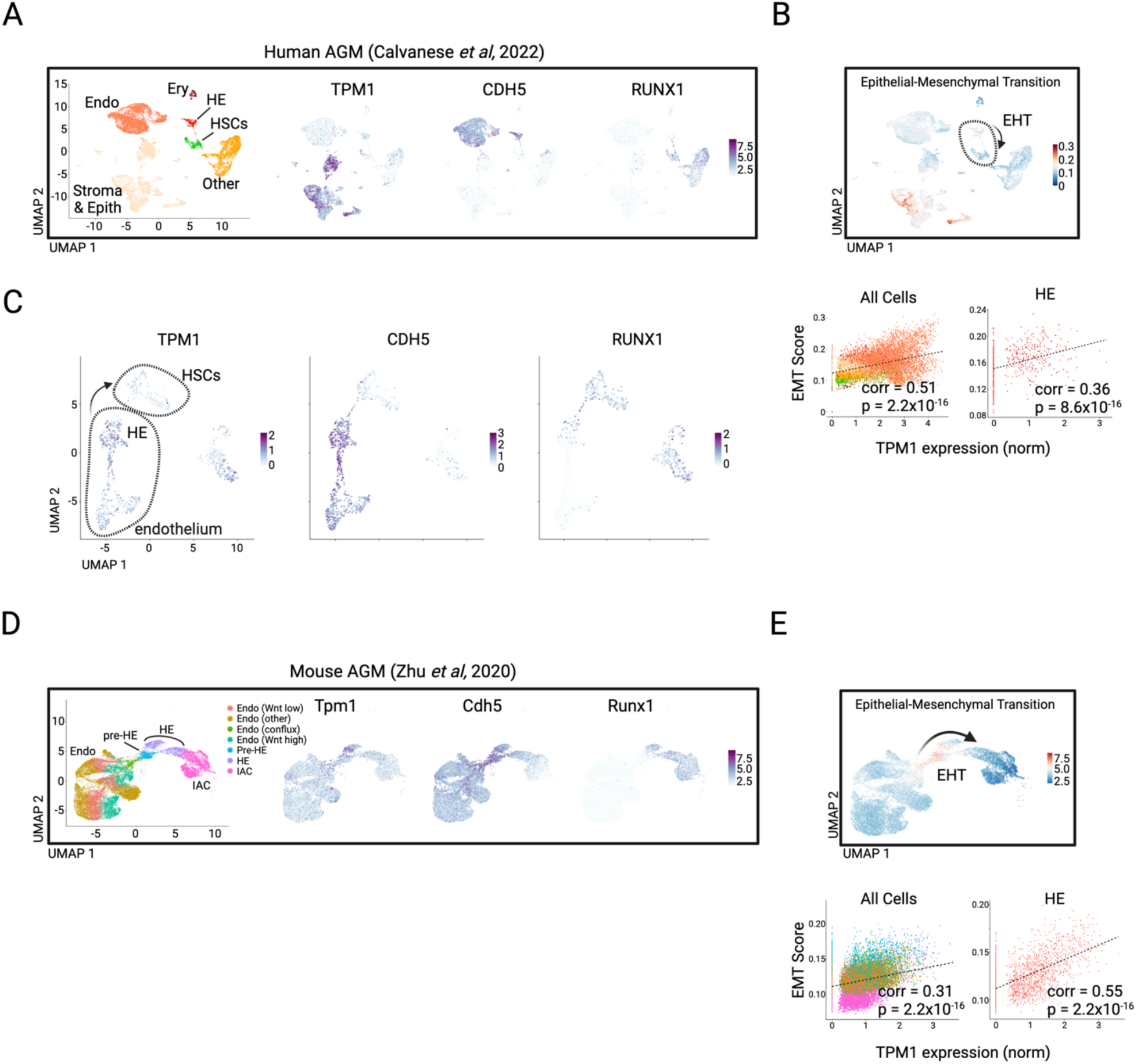
Single cell analyses link *TPM1* expression and EMT/EHT during definitive human and murine hematopoiesis. (A) Single cell RNA sequencing analysis of CD34^+^ and/or CD31^+^ enriched cells from human AGM (CS14-15) ^32^, with UMAP plots highlighting *TPM1*, *CDH5* (*VE-CAD*), and *RUNX1* expression. Cell identity labels are based on gene expression patterns from Calvanese *et al* ^32^. (B) EMT scores for human AGM cells based on expression of human Hallmark Epithelial-to-Mesenchymal Pathway genes ^34^. Red color specifies cells with higher normalized expression of EMT genes. Scatterplots depict Pearson correlation and related p-values for normalized *TPM1* expression and EMT scores for all analyzed cells or HE cells. (C) UMAP analysis of hemato-vascular cell populations from human AGM, highlighting *TPM1* expression during EHT. *CDH5* and *RUNX1* serve as HE and HPC emergence landmarks. Arrow depicts EHT, with gatings as depicted in Calvanese *et al* ^32^. (D) Single cell sequencing analysis of murine cells derived from AGM, highlighting the EHT, with UMAP plots highlighting *Tpm1*, *Cdh5* (*VE-Cad*), and *Runx1* expression. Cell cluster labels are based on Zhu *et al* ^33^. (E) EMT scores for murine AGM cells based on expression of murine Hallmark Epithelial-to-Mesenchymal Pathway genes ^34^. Scatterplots depict Pearson correlation and related p-values for normalized *TPM1* expression and EMT scores for all analyzed cells or HE cells.

Interestingly, *TPM1* expression correlated with EMT/EHT progression in this data set. When we scored each cell based on its expression of established EMT-related genes ^34^, we noted that higher EMT scores correlated with increased *TPM1* expression (**Fig. 4B**). Targeted analysis of cells undergoing EHT revealed that *TPM1* expression increased slightly in HE and was downregulated in HPCs following EHT completion (**Fig. 4B-C**). The observed increase in *TPM1* expression in cells undergoing EMT was also consistent with prior data from other cellular systems ^21–25^, and these observations extended to the murine hematopoietic system. Similar to human development, murine Runx1^+^Cdh5^+^ HE form Runx1^+^ HPCs that ultimately support lifelong hematopoiesis ^33^. Our analysis of single-cell transcriptional profiles showed that *Tpm1* was expressed at increased levels in murine pre-hemogenic endothelium (pre-HE) and HE cells, with subsequent downregulation in HPCs (**Fig. 4D**). Similar to human hematopoiesis, murine cells undergoing EHT exhibited increased *Tpm1* expression (**Fig. 4E**).

### Tropomyosin 1 deficiency enhances murine HE specification

We hypothesized that *Tpm1* may have the same impact on murine hematopoiesis as we observed in the human in vitro system. We obtained a *Tpm1* GeneTrap-Reporter mouse model (*Tpm1^GT^*) ^35–38^, which contains an intronic splice acceptor site linked to a β-galactosidase reporter gene (LacZ) positioned to capture all *Tpm1* isoform transcripts (**Supplementary Fig. 5S1A**). We confirmed that this construct efficiently captured *Tpm1* transcripts by observing that *Tpm1^GT/+^* mice had decreased Tpm1 protein in their peripheral blood, and by observing embryonic lethality in *Tpm1^GT/GT^* embryos (**Supplementary Fig. 5S1B-C**). The timing of embryonic lethality was consistent with other *Tpm1* knockout mouse models, which have reported severe cardiac dysmorphology ^39^. The presence of LacZ reporter expression in the E9.5 dorsal aortic endothelium was also consistent with our expectations for *Tpm1* expression, based on protein expression during in vitro hematopoiesis (**Supplementary Fig. 5S1D** vs **Fig. 1B**).

We used the *Tpm1^GT^* mouse model to determine if *Tpm1* deficiency enhanced HE specification and HPC formation in vivo. During murine development, morphologically flat Runx1^+^CD31^+^ HE appear around embryonic day 9.5 post-conception (E9.5) and subsequently form round cKit^+^Runx1^+^ intra-aortic clusters (IACs) of HPCs around E10.5 ^33,40^ (**Fig. 5A-B**). By whole-mount imaging ^40^, *Tpm1^GT/+^* embryos had enhanced quantities of HE cells at E9.5 and IACs at E10.5 (**Fig. 5B-C**). The number of HE cells was normal at E10.5, as was the quantity of IACs at E11.5 (**Fig. 5C**). These findings suggested that *TPM1* haploinsufficiency increased HE and HPC production between E9.5-E10.5 in vivo. We then used limiting dilution assays ^33^ to quantify B or T lymphoid progenitor cell frequencies in E10.5 *TPM1^GT/+^*embryos. There was a trend toward increased lymphoid progenitor frequencies in *TPM1^GT/+^* embryos compared with littermate controls (∼17% increase), although these data did not meet statistical significance due to variation across embryos (**Supplementary Fig. 5S2**). Taken together, these findings show that *Tpm1* deficiency enhanced HE specification without compromising HPC function.

**Figure 5.**
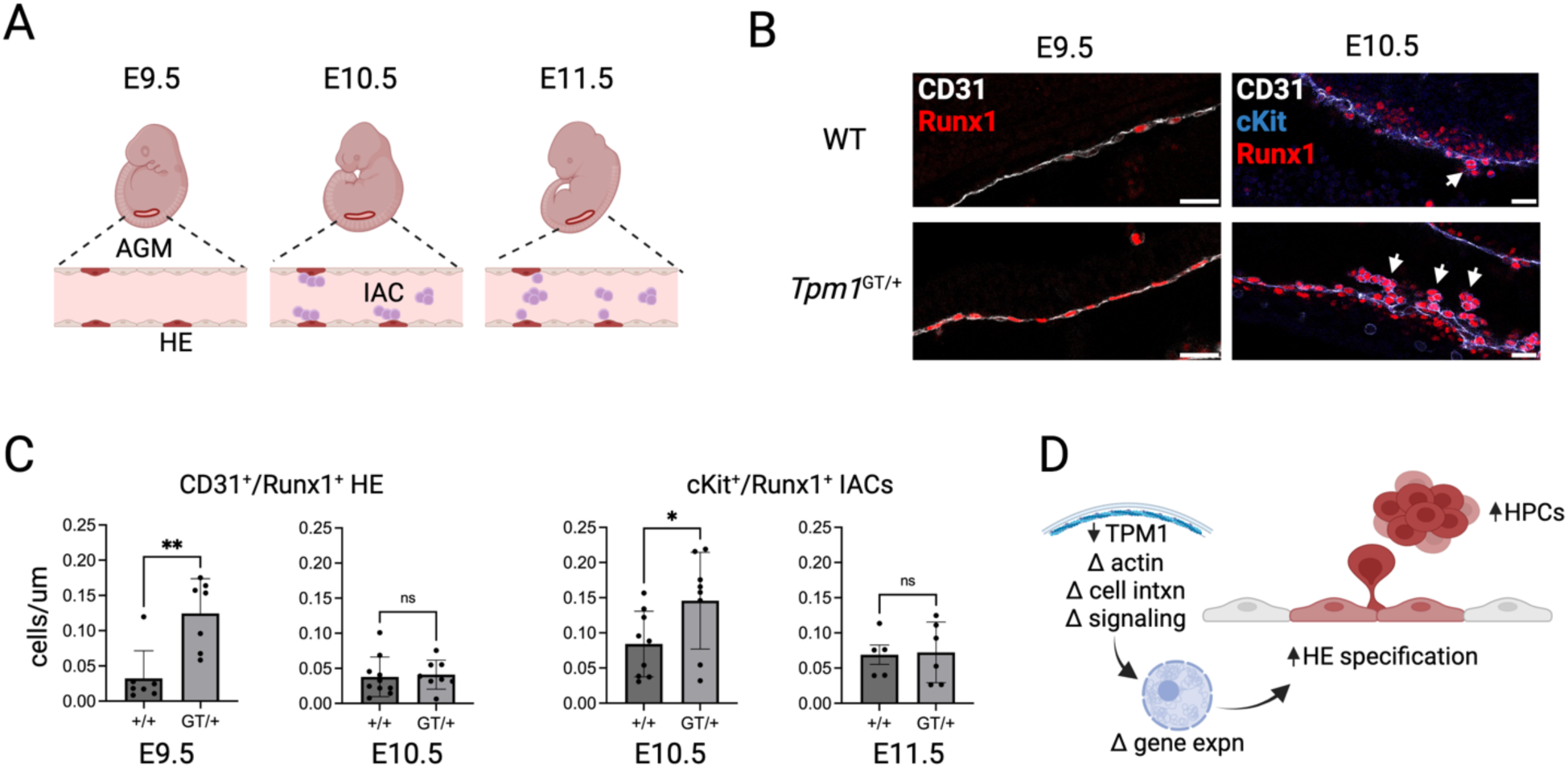
*Tropomyosin 1* regulates HE and HPC formation during in vivo hematopoiesis. (A) During murine embryogenesis, CD31^+^Runx1^+^ HE cells in the major caudal arteries emerge around E9.5. These HE cells form intra-aortic clusters (IACs) of cKit^+^Runx1^+^ HPCs on E10.5-E11.5, which will seed fetal liver and later migrate to the bone marrow. (B) Representative whole mount staining of E9.5 and E10.5 aorta-gonad-mesonephros (AGM) from WT and *Tpm1^GT/+^* embryos. Arrows point to IACs. Scale bars, 50uM. (C) Quantifications of imaging studies show that heterozygous *Tpm1^GT/+^* mice have increased frequencies of HE cells at E9.5 and IAC cells at E10.5 compared to littermate controls. Visualizations were centered on the intersection of vitelline artery and dorsal aorta. *p<0.05. **p<0.01 by t test. (D) Actin cytoskeletal alterations in the context of *Tpm1* deficiency promote gene expression changes that HE cell specification during hematopoietic development.

## Discussion

This study reveals a novel role for *TPM1* in developmental hematopoiesis across species and hematopoietic ontogeny ^11,19^, from in vitro human primitive hematopoiesis to in vivo murine definitive hematopoiesis. *TPM1* downregulation normally occurs during HE specification in human cells, and *TPM1* deficiency is sufficient to augment the formation of HE and functional HPCs. Many studies have focused on transcription factors (e.g., *Runx1*) ^19,41^ that drive hematopoiesis through ‘inside-out’ mechanisms – i.e., transcriptional changes that promote altered morphology and cellular development. *TPM1*-related mechanisms instead represent a distinct ‘outside-in’ paradigm wherein altered actin cytoskeletal dynamics and cell interactions impact downstream transcriptional and/or developmental programs (**Fig. 5D**). Indeed, actomyosin contractility is active during hematopoiesis ^27^ and biomechanical forces from blood flow promote blood cell formation ^42,43^. Since *TPM1* is normally highly expressed in stromal and arterial endothelial populations, *TPM1* deficiency may also promote HE specification by inhibiting vascular endothelial programming, an alternative fate during endothelial development. Future studies will determine if these changes occur via direct impacts on developing HE precursors, or indirectly through alterations in the *Tpm1*-deficient stromal environment.

Mammalian cells can express >40 tropomyosin isoforms and *TPM1* isoforms can have dramatically different biological effects ^5^. Our findings indicate that the aggregated effect of *TPM1* gene products is to constrain, but not compromise, hematopoiesis. A limitation of this study is that all in vitro and in vivo models presented herein reflect coordinate depletion of all *TPM1* isoforms. While future studies are needed to conditionally delete specific *Tpm1* isoforms, high molecular weight *TPM1* isoforms (e.g., *TPM1.6/1.7*) seem most likely to directly constrain HE specification and/or EHT, given established effects in cell models ^21–25^ and an increased presence of these isoforms in adherent cell types (**Fig. 1B**). However, it has thus far been difficult to define functions for specific *TPM1* isoforms using overexpression studies during in vitro hematopoiesis (data not shown). Multiple isoforms may contribute independently to the regulatory effect of *TPM1* on normal hematopoiesis.

Effects observed herein may also have relevance for non-hematopoietic tissue development. Increased *TPM1* expression in stromal and epithelial cells undergoing EMT may indicates a more general regulatory role for *TPM1* during cell state transitions in embryonic cells (**Fig. 4**). This is consistent with previous studies that have also suggested links between TPM1 and EMT, although the role of TPM1 in promoting or inhibiting EMT progression has differed depending on cell or tissue context. Whereas concurrent *TPM1* and *TPM2* knockout have a deleterious effect on ocular lens development ^6^, *TPM1* deficiency seems to promote EMT in cancer models ^21–25^. The hematopoietic system represents another example wherein *TPM1* deficiency positively impacts an EMT-like process.

In vitro-derived blood cells have recently been shown to support the production of clinical testing reagents ^44^ and cell therapeutics ^45–47^, but blood cell yields remain inefficient. Factors influencing HE specification have been elusive, but could be co-opted to enhance in vitro hematopoiesis.

Our findings suggest that temporal modulation of TPM1 may represent a novel strategy to augment HE formation. The general cellular mechanisms by which *TPM1* regulates endothelial cell production during in vitro hematopoiesis may also facilitate derivation of other endothelial populations, including production of pulmonary endothelial cells to support cellular therapeutics development ^48,49^.

In addition to defining a novel role for *TPM1* in hematopoiesis and perhaps hinting at mechanisms responsible for genetic associations linking polymorphisms in the *TPM1* gene locus with altered human blood traits ^8,9^, our findings raise interesting questions about the regulation of different hematopoietic progenitor populations. Multiple waves of HE cells produce HPCs with different engraftment and differentiation potentials ^32,50^. Future studies are needed to define which specific subset(s) of HE cells are affected by *TPM1-*mediated actin regulation.

More broadly, the radical changes required to form HE and HPCs during hematopoiesis represent an exciting area of study for actin and tropomyosin biology.

## Methods

### Stem cell culture and differentiation

Wild type control iPSC lines were obtained from the CHOP Pluripotent Stem Cell Core. *TPM1* KO cell lines were created using targeted CRISPR/Cas9 ^51^ and subsequently validated ^10,28^. iPSCs were maintained on mouse embryonic fibroblast feeders until differentiation. Prior to differentiation, iPSC were cultured in feeder-free conditions for at least 2 passages.

### Western blots

Collected cell pellets were lysed in RIPA and separated by electrophoresis on 4-12% NuPAGE Bis-Tris Mini Protein gels (Invitrogen), followed by semi-dry transfer to nitrocellulose membranes using a Power Blotter apparatus (Invitrogen). After blocking in 3% milk in TBST, membranes were sequentially incubated with primary and secondary antibodies. Washed blots were imaged using a ChemiDoc (BioRad) or c-Digit (LiCor) apparatus. Antibodies used for these studies are listed in **Supplementary Table 5**.

### Flow cytometry

Flow cytometry experiments utilized a Cytoflex LX (Beckman Coulter). Data were analyzed and presented using FlowJo software (BD Biosciences). Antibodies used for these studies are listed in **Supplementary Table 5**.

### Cell proliferation analysis

Cell proliferation utilized CFSE staining per manufacturer’s instructions (ThermoFisher Scientific). Prewarmed media containing CFSE stain was added to cells on day 0, and fluorescence intensity measured by flow cytometry at intervals thereafter. As an alternative marker of cell proliferation and additional quality control, harvested cells were manually counted using a CytoSMART hemacytometer (Axion Biosystems).

### Cell cycle analysis

EdU staining assay was performed according the manufacturer’s instructions (ThermoFisher Scientific). At the time points indicated in the text, cells were cultured at 37°C in EdU stain for 2 h prior to harvest. Collected cells were washed with 1% BSA in PBS, fixed, and stained with Click-iT™ reagents per kit instructions along with FxCycle™ Violet Stain to mark DNA content (ThermoFisher Scientific) and developmental stage-related cell surface markers. Cells were analyzed by flow cytometry on a Cytoflex LX instrument (Beckman Coulter).

### Bulk RNA sequencing

Bulk RNA was isolated from iPSCs after feeder-free passage (day 0) and iPSC-derived cells, including mesoderm (day 2 cultures), FACS-sorted KDR^+^CD31^+^ endothelial cells (day 4), or non-adherent HPCs (day 8-9), using a PureLink RNA Micro kit (Invitrogen) according the manufacturer’s instructions. CHOPWT14 and CHOP14 *TPM1KO* iPSC, endothelium, and HPCs libraries were prepared using a Takara Total RNA with ribosomal depletion kit. All other sample libraries were prepared using an Illumina TruSeq stranded kit. We constrained direct comparisons to samples prepared using analogous library preparation. Raw data were aligned using STAR (v2.7.10) and analyzed using DEBrowser (v1.24.1, ^52^). Presented results were obtained using DESeq2 after RLE normalization, with significant differential gene expression set at p<0.05 and fold change>1.5. Differentially expressed gene lists were then analyzed using Enrichr ^53–55^.

### Limiting dilution assay quantitation

Endothelial cells (CD31^+^ or CD34^+^) were isolated from primitive hematopoietic differentiations and sorted on a MoFlo Astrios (Beckman Coulter) into pre-treated 96-well tissue culture plates containing growth factor-reduced Matrigel. Cells were plated at 3-1000 cells per well and cultured for 1 week in bFGF, SCF, Flt3L, and VEGF. Media was added every 2-3 days. After a week, adherent and non-adherent cells were harvested from each well, stained, and analyzed by flow cytometry. Wells containing at least 10 CD43^+^ HPCs were deemed ‘positive’ as having initially contained HE. HE frequency was calculated using extreme limiting dilution analysis (ELDA) software ^56^.

For murine embryo limiting dilution assays, whole E10.5 embryos were dissociated and processed as described ^33^. Lymphocyte progenitor frequencies were calculated using ELDA software ^56^.

### Colony formation assays

Colony assays were performed according to manufacturer’s instructions with MegaCult^TM^-C or Methocult^TM^ H4435 Enriched media (StemCell Technologies) using fresh or cryopreserved HPCs. For Methocult assays, cells were cultured in a 35 mm dish and manually counted after 12-14 days. For MegaCult assays, cells were cultured, fixed after 10-12 days, and manually counted. Colony counts were performed on an Olympus IX70 microscope and colony images acquired with a Zeiss Axiocam 208 color camera.

### Single cell RNA sequencing analysis

Publicly available single cell RNA seq data were acquired for human ^32^ or mouse ^33^. For human data, we integrated CS14-CS15 data sets using the R package Harmony ^57^ and annotated cell types using Seurat (v4.3.0) according to previously described cell surface markers ^32^. We used the R package UCell ^58^ to calculate EMT scores, according to genes specified in the MSigDB Hallmark EMT gene pathway ^34^. Pearson coefficients, calculated to correlate EMT scores with *TPM1* expression, are presented in each scatterplot.

For mouse scRNAseq analysis, primary data ^33^ were integrated using Harmony ^57^ and analyzed for select gene expression. We used UCell ^58^ to calculate EMT scores after identifying murine analogs for genes in the MSigDB Hallmark EMT gene pathway ^34^. EMT scores were correlated with *TPM1* expression based on the Pearson coefficients shown in each scatterplot.

### Mouse line derivation

All mouse studies were approved by the Children’s Hospital Institutional Animal Use and Care Committee. The *TPM1* GeneTrap-Reporter mouse construct was initially rederived from cryopreserved embryos with the assistance of the CHOP Transgenic Core Facility. We thank the Wellcome Trust Sanger Institute Mouse Genetics Project (Sanger MGP) and its funders for providing the mutant mouse line (129-TPM1<tm1a(EUCOMM)Wtsi>/WtsiH), and INFRAFRONTIER/EMMA (www.infrafrontier.eu) partner MRC Harwell from which the mouse line was received. Funding information may be found at www.sanger.ac.uk/mouseportal and associated primary phenotypic information at www.mousephenotype.org ^35–38^. These mice were bred and backcrossed to C57Bl6/J mice (Jackson Labs) for at least 2 generations prior to analysis.

### Whole mount embryo imaging

Whole mount imaging was carried out as previously described ^40^. Embryos were harvested at the indicated time points after timed matings and fixed in methanol prior to staining with the antibodies indicated in **Supplementary Table 5**. After washing, embryos were mounted on glass slides and imaged using a Leica SP8 confocal microscope. Visualizations were centered on the intersection of the vitelline artery and dorsal aorta. HE and IACs were manually identified and quantified from resultant images using ImageJ software ^59^.

### Statistics and data plotting

Statistics and data were calculated and plotted using GraphPad Prism 9 or R (v4.2.2). Graphical schematics were generated using BioRender (www.BioRender.com).

### Code and data availability

All coding scripts and data are available by request. Public single cell RNA sequencing analyses were collected from GSE137117 (murine) and GSE162950 (human).

## Acknowledgements

We thank the Children’s Hospital of Philadelphia (CHOP) Flow Cytometry, Pluripotent Stem Cell, and Mouse Transgenic Cores for their assistance with this study. We thank Drs. Harry Ischiropoulos, Serena Raimo, and Ipsita Mohanty for their helpful guidance and assistance. We thank John Daniels, Nasir Riley, and the CHOP High Performance Computing Cluster support team for their help throughout this project.

## Supplementary Information

### Supplemental Figures

**Fig 1 S1.**
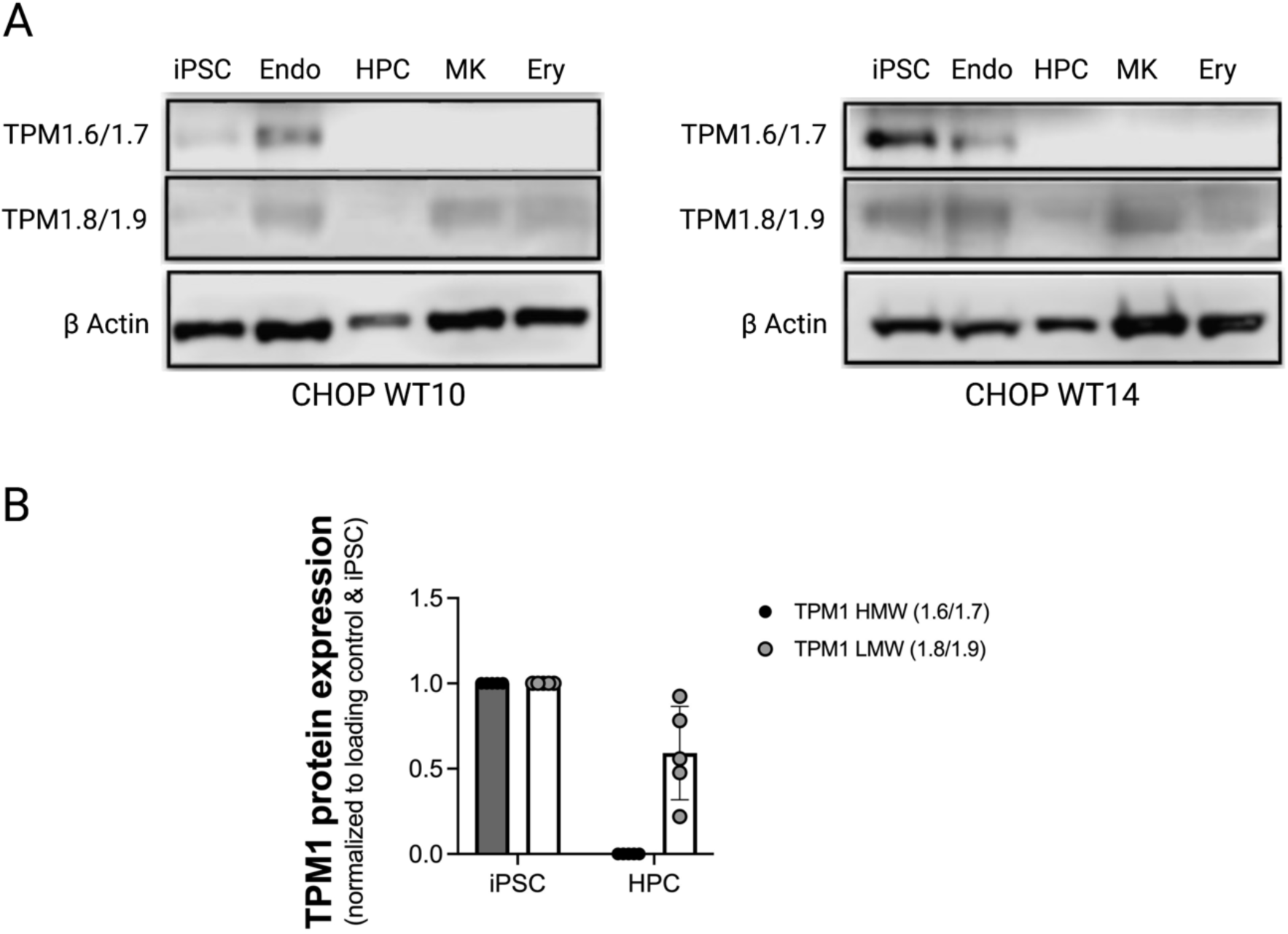
TPM1 isoform presence during hematopoietic development. (A) High molecular weight TPM1.6/1.7 isoforms are expressed at the adherent (iPSC and endothelial) cell stages but not in non-adherent primitive in vitro-derived HPCs, megakaryocytes (MK) and erythroid (Ery) cells. Low molecular weight TPM1.8/1.9 isoforms remain present throughout in vitro differentiation in adherent and non-adherent cell types. Left and right panels are derived from 2 separate wild type (WT) iPSC lines. (B) Protein quantifications from western blots from these isogenic lines (n=5 blots total from two WT iPSC lines). TPM1 isoform presence in HPCs was normalized to loading control and plotted relative to iPSC value.

**Fig 1 S2.**
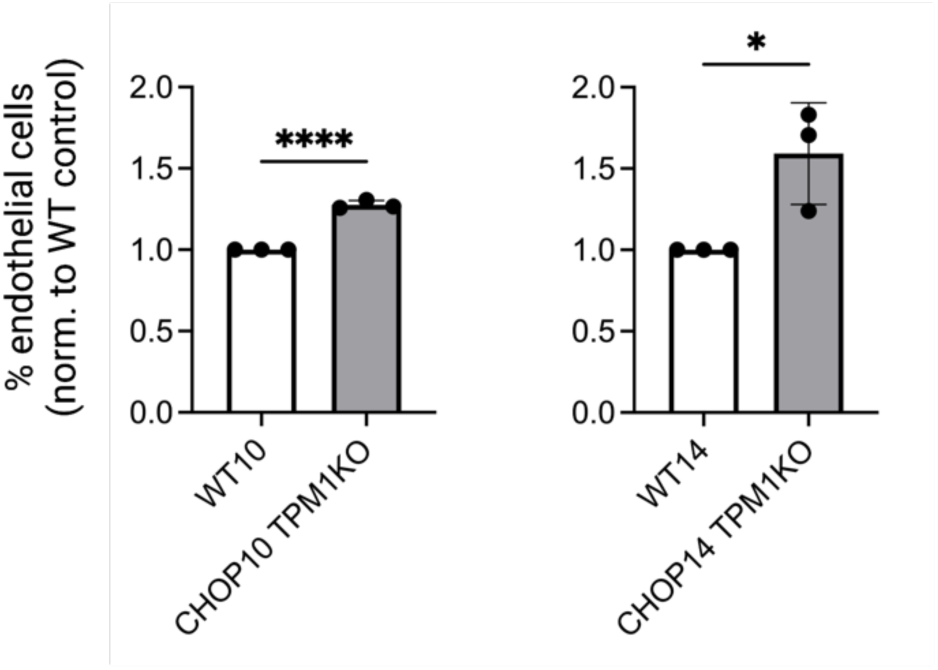
*TPM1KO* increases endothelial cell production during in vitro hematopoiesis in the CHOP WT10 and CHOP WT14 backgrounds. In a CHOP WT10 background or CHOP WT14 background, cells containing a genomic deletion of *TPM1* produce more endothelial cells (CD31^+^ and/or CD34^+^) during primitive hematopoietic differentiation (n=3 experiments per line).

**Fig 1 S3:**
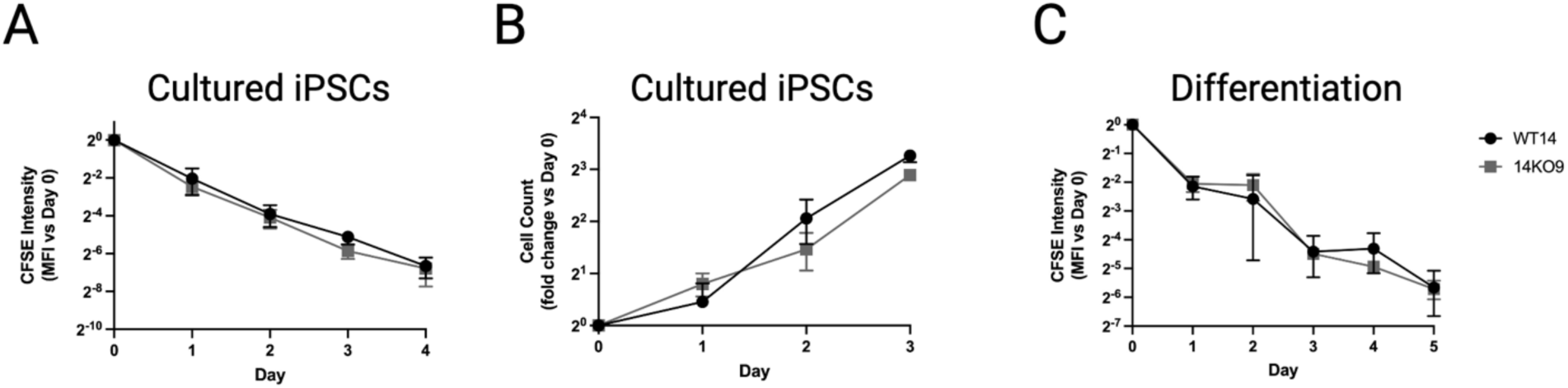
*TPM1KO* does not impact cell proliferation in a CHOP WT14 genetic background. (A) CFSE staining intensity washout does not significantly differ in WT and *TPM1KO* iPSCs. (B) Direct cell counts do not significantly differ between WT and *TPM1KO* iPSCs (n=4 experiments). (C) CFSE staining intensity washout does not significantly differ between WT and *TPM1KO* cells during in vitro differentiation (n=5 differentiations).

**Fig 1 S4.**
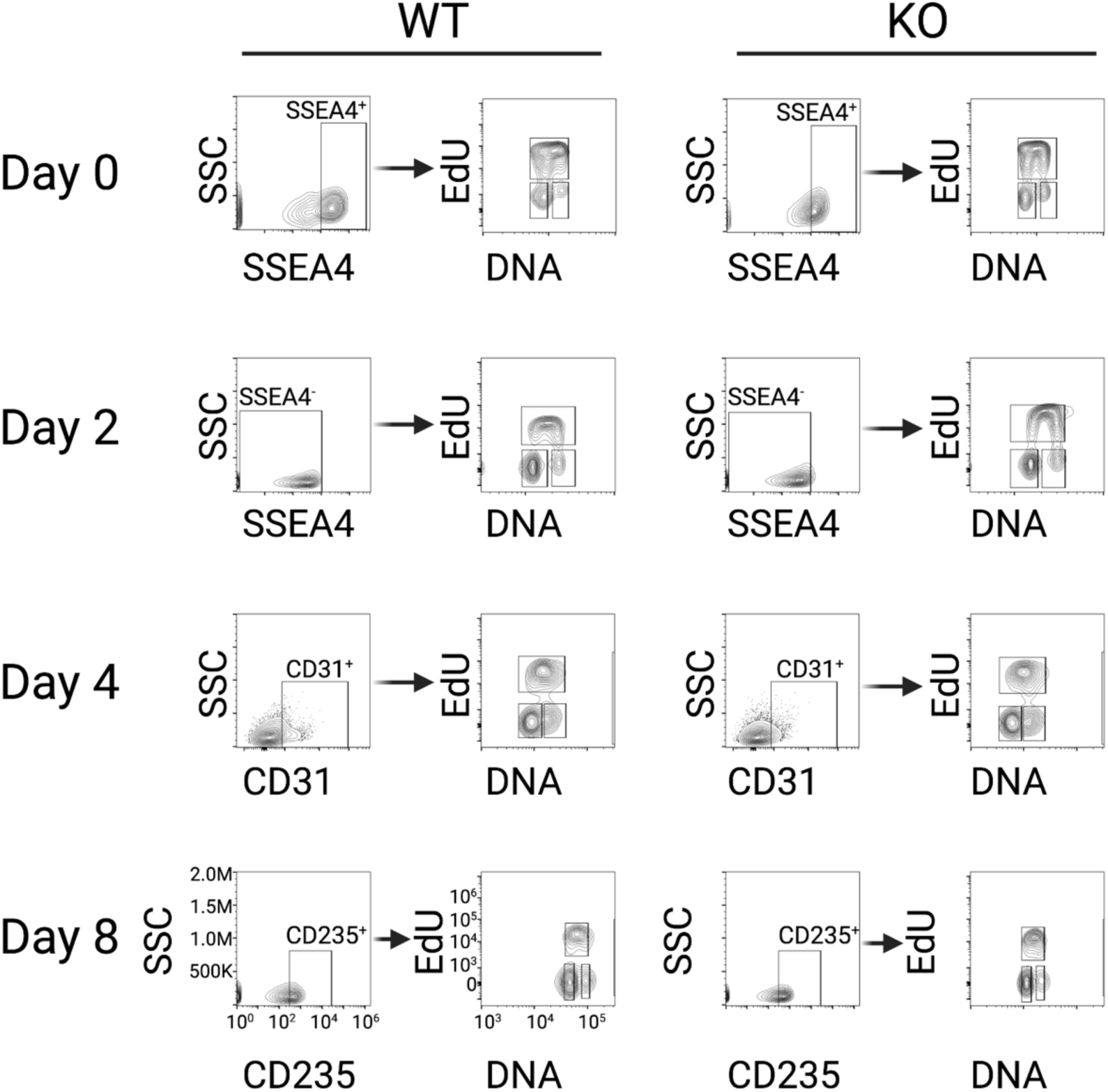
EdU flow cytometry gating strategy for cell cycle analysis. iPSCs, mesoderm (day 2 SSEA4^-^ cells), CD31^+^ endothelial, and CD235^+^ hematopoietic progenitor cells were identified. Some experiments analyzed all iPSCs, as we did not note differences when analyzing SSEA4^+^ iPSCs or total cultures. At each time point, the proportion of cells in G0/G1, S, or G2/M phases of the cell cycle were quantified.

**Fig 2 S1:**
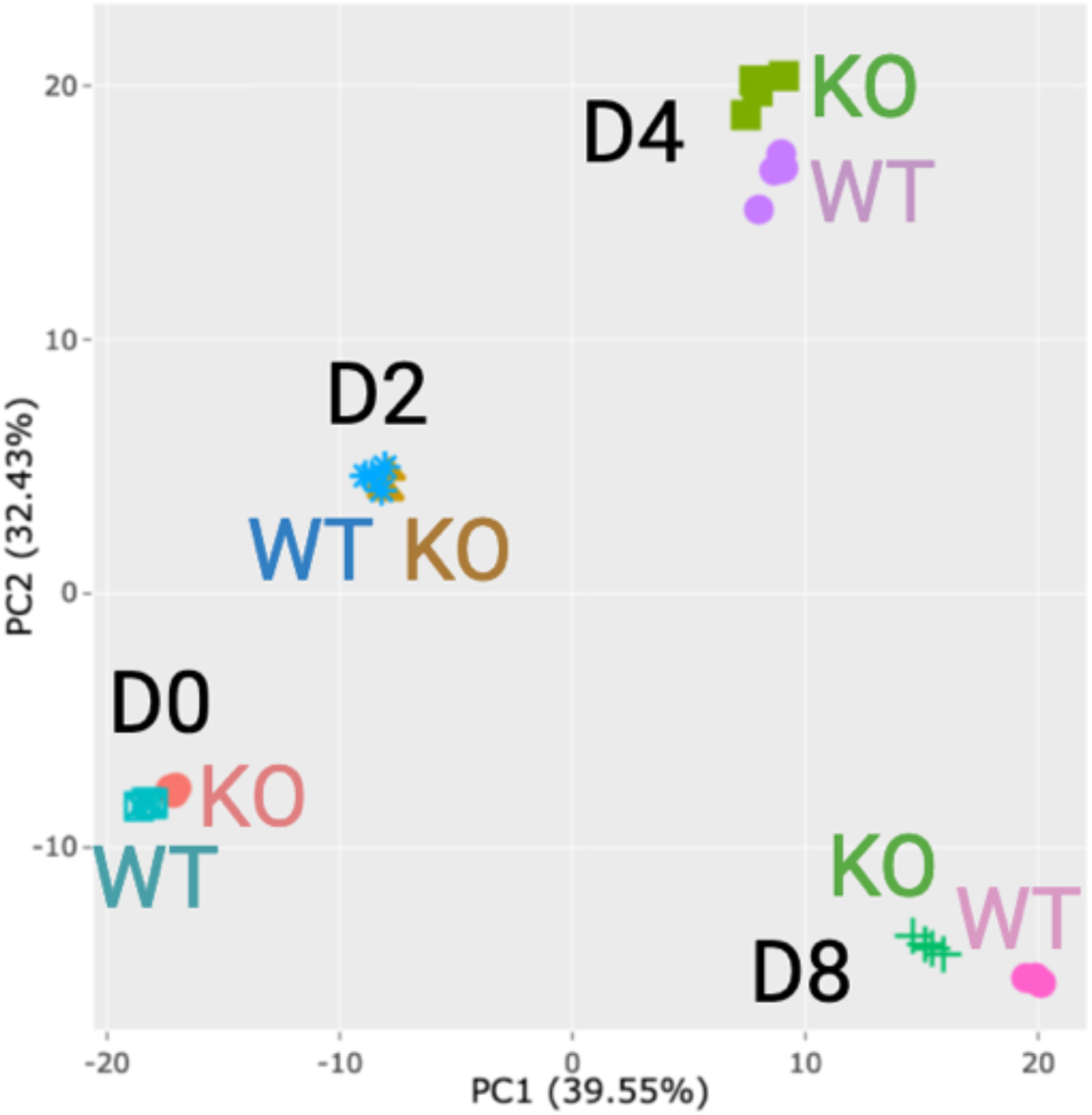
Bulk RNA sequencing transcriptomic analyses show that *TPM1KO* does not globally disrupt gene expression systems during primitive hematopoietic differentiation. PCA plot shows that *TPM1KO* and isogenic CHOP WT14 controls are similar throughout differentiation, including on day 0 (iPSC), day 2 (mesoderm), day 4 (endothelial), and day 8 (HPC) cell stages.

**Fig 3 S1.**
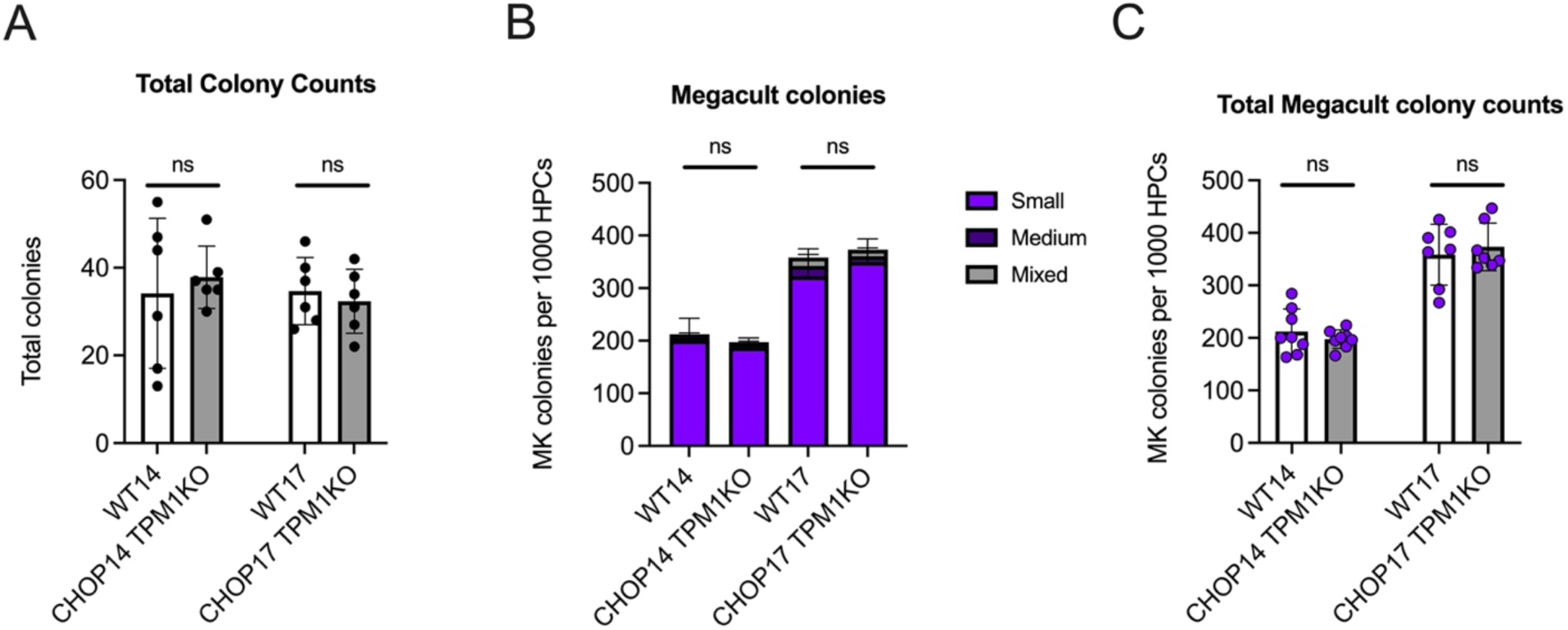
*TPM1KO* does not alter primitive HPC lineage production. (A) Total colony counts (erythroid, myeloid, mixed) did not differ between *TPM1KO* and WT controls across 2 isogenic backgrounds (WT14 and WT17, n=6 assays per genotype). (B-C) Megakaryocyte colony production did not significantly differ between *TPM1KO* and WT controls across 2 isogenic backgrounds (WT14 and WT17). This was true in terms of (B) colony size and cell composition and (C) total megakaryocyte colony counts (n=7-8 assays per group).

**Fig 5 S1.**
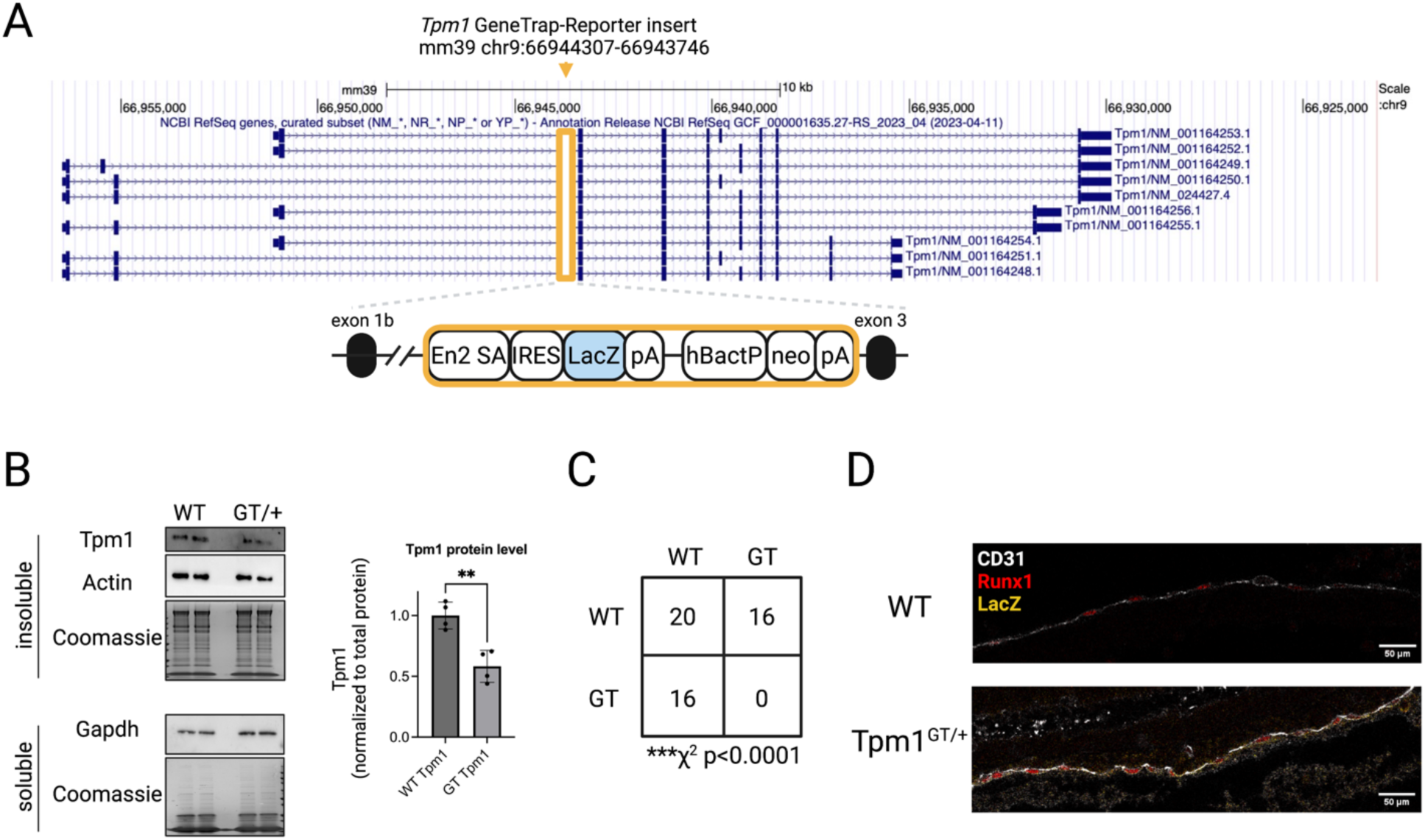
*Tpm1* GeneTrap-Reporter mouse model validation. (A) Construct map detailing insertion point of En2 Splice Acceptor (SA)-IRES-LacZ-polyA and neomycin resistance at the indicated position in the murine mouse genome. Note that in the gene construct map showing different *Tpm1* isoforms, transcription proceeds right to left. (B) By western blot, low molecular weight Tpm1 protein level (Tpm1.8/1.9) is reduced ∼50% in the insoluble (cytoskeletal and membrane) fraction of *Tpm1^GT/+^* murine whole blood. (C) Heterozygous *Tpm1^GT/+^*x *Tpm1^GT/+^* breedings did not produce viable *Tpm1^GT/GT^* offspring (p<0.0001 by Chi-square test), indicating embryonic lethality. Numbers indicate pup genotypes that contributed to this analysis. (D) In E9.5 embryos, *Tpm1^GT/+^*but not wild type littermate controls express LacZ in the aorta-gonad-mesonephros (AGM) region that produces Runx1^+^CD31^+^ hemogenic endothelium. Scale bars, 50 μm.

**Fig 5 S2.**
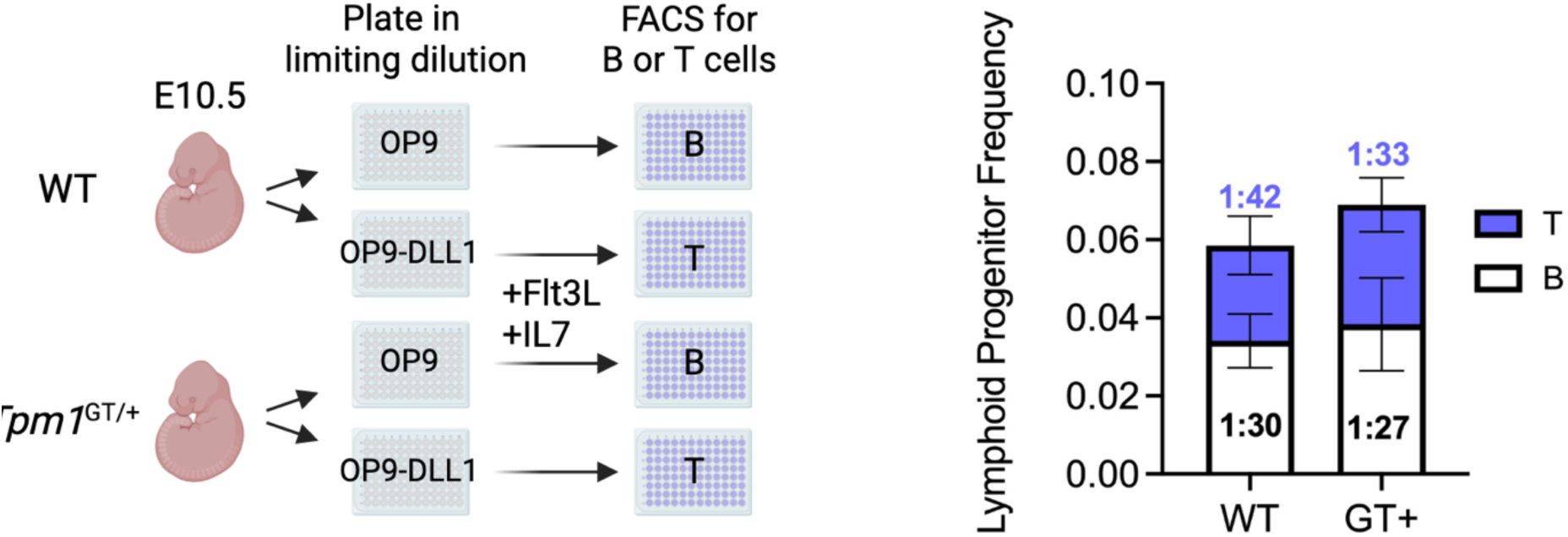
*Tpm1^GT/+^* does not significantly increase E10.5 AGM lymphoid progenitor frequency. AGM regions from E10.5 embryos were dissociated and plated in limiting dilution on OP9 or OP9-DLL1 cells to culture B or T cells, respectively. Bar plot represents estimated lymphoid progenitor cell frequencies. Mean frequencies for B and T progenitors are indicated. Bars represent mean with 95% confidence intervals (n=6-9 embryos per group).

## References

1. Meiring, J. C. M. et al. Co-polymers of Actin and Tropomyosin Account for a Major Fraction of the Human Actin Cytoskeleton. Current Biology 28, 2331–2337.e5 (2018).

2. Gateva, G. et al. Tropomyosin Isoforms Specify Functionally Distinct Actin Filament Populations In Vitro. Current Biology 27, 705–713 (2017).

3. Schevzov, G., Whittaker, S. P., Fath, T., Lin, J. J.-C. C. & Gunning, P. W. Tropomyosin isoforms and reagents. Bioarchitecture 1, 135–164 (2011).

4. Brayford, S. et al. Tropomyosin promotes lamellipodial persistence by collaborating with Arp2/3 at the leading edge. Current Biology 26, 1312–1318 (2016).

5. Gunning, P. W. & Hardeman, E. C. Tropomyosins. Current Biology 27, R8–R13 (2017).

6. Shibata, T. et al. Lens-specific conditional knockout of tropomyosin 1 gene in mice causes abnormal fiber differentiation and lens opacity. (2021) doi:10.1016/j.mad.2021.111492.

7. Kubo, E., Hasanova, N., Fatma, N., Sasaki, H. & Singh, D. P. Elevated tropomyosin expression is associated with epithelial-mesenchymal transition of lens epithelial cells. J Cell Mol Med 17, 212–221 (2013).

8. Chen, M. H. et al. Trans-ethnic and Ancestry-Specific Blood-Cell Genetics in 746,667 Individuals from 5 Global Populations. Cell 182, 1198–1213.e14 (2020).

9. Vuckovic, D. et al. The Polygenic and Monogenic Basis of Blood Traits and Diseases. Cell 182, 1214–1231.e11 (2020).

10. Thom, C. S. et al. Tropomyosin 1 genetically constrains in vitro hematopoiesis. BMC Biol 18, 52 (2020).

11. Dzierzak, E. & Speck, N. A. Of lineage and legacy: The development of mammalian hematopoietic stem cells. Nature Immunology vol. 9 129–136 Preprint at 10.1038/ni1560 (2008).

12. Lucitti, J. L. et al. Vascular remodeling of the mouse yolk sac requires hemodynamic force. Development 134, 3317–3326 (2007).

13. Goldie, L. C., Lucitti, J. L., Dickinson, M. E. & Hirschi, K. K. Cell signaling directing the formation and function of hemogenic endothelium during murine embryogenesis. Blood 112, 3194–3204 (2008).

14. Gritz, E. & Hirschi, K. K. Specification and function of hemogenic endothelium during embryogenesis. Cellular and Molecular Life Sciences 73, 1547–1567 (2016).

15. Marcelo, K. L. et al. Hemogenic endothelial cell specification requires c-Kit, notch signaling, and p27-mediated cell-cycle control. Dev Cell 27, 504–515 (2013).

16. Coulombe, P. et al. Meis1 establishes the pre-hemogenic endothelial state prior to Runx1 expression. Nat Commun 14, 4537 (2023).

17. Espín-Palazón, R. et al. Proinflammatory signaling regulates hematopoietic stem cell emergence. Cell 159, 1070–1085 (2014).

18. Howell, E. D. et al. Efficient hemogenic endothelial cell specification by RUNX1 is dependent on baseline chromatin accessibility of RUNX1-regulated TGFβ target genes. Genes Dev 35, 1475–1489 (2021).

19. Ottersbach, K. Endothelial-to-haematopoietic transition: An update on the process of making blood. Biochemical Society Transactions vol. 47 591–601 Preprint at 10.1042/BST20180320 (2019).

20. Kalluri, R. & Weinberg, R. A. The basics of epithelial-mesenchymal transition. Journal of Clinical Investigation vol. 119 1420–1428 Preprint at 10.1172/JCI39104 (2009).

21. Wang, J., Tang, C., Yang, C., Zheng, Q. & Hou, Y. Tropomyosin-1 functions as a tumor suppressor with respect to cell proliferation, angiogenesis and metastasis in renal cell carcinoma. J Cancer 10, 2220–2228 (2019).

22. Bakin, A., Varga, A., Zheng, Q. & Safina, A. Epigenetic silencing of tropomyosin alters transforming growth factor beta control of cell invasion and metastasis. Breast Cancer Research 7, P4.15 (2005).

23. Bakin, A. V et al. A Critical Role of Tropomyosins in TGF-Regulation of the Actin Cytoskeleton and Cell Motility in Epithelial Cells. Mol Biol Cell 15, 4682–4694 (2004).

24. Dai, Y. & Gao, X. Inhibition of cancer cell-derived exosomal microRNA-183 suppresses cell growth and metastasis in prostate cancer by upregulating TPM1. Cancer Cell Int 21, (2021).

25. Pan, H. et al. Tropomyosin-1 acts as a potential tumor suppressor in human oral squamous cell carcinoma. PLoS One 12, 1–13 (2017).

26. Gagat, M. et al. CRISPR-Based Activation of Endogenous Expression of TPM1 Inhibits Inflammatory Response of Primary Human Coronary Artery Endothelial and Smooth Muscle Cells Induced by Recombinant Human Tumor Necrosis Factor α. Front Cell Dev Biol 9, (2021).

27. Lancino, M. et al. Anisotropic organization of circumferential actomyosin characterizes hematopoietic stem cells emergence in the zebrafish. Elife 7, (2018).

28. Wilken, M. B. et al. Generation of a human Tropomyosin 1 knockout iPSC line. bioRxiv (2023) doi:10.1101/2023.05.03.539242.

29. Canu, G. et al. Analysis of endothelial-to-haematopoietic transition at the single cell level identifies cell cycle regulation as a driver of differentiation. Genome Biol 21, 157 (2020).

30. Colin, A., Bonnemay, L., Gayrard, C., Gautier, J. & Gueroui, Z. Triggering signaling pathways using F-actin self-organization. Sci Rep 6, (2016).

31. Saxena, S., Rönn, R. E., Guibentif, C., Moraghebi, R. & Woods, N. B. Cyclic AMP Signaling through Epac Axis Modulates Human Hemogenic Endothelium and Enhances Hematopoietic Cell Generation. Stem Cell Reports 6, 692–703 (2016).

32. Calvanese, V. et al. Mapping human haematopoietic stem cells from haemogenic endothelium to birth. Nature 604, 534–540 (2022).

33. Zhu, Q. et al. Developmental trajectory of prehematopoietic stem cell formation from endothelium. Blood 136, 845–856 (2020).

34. Liberzon, A. et al. The Molecular Signatures Database Hallmark Gene Set Collection. Cell Syst 1, 417–425 (2015).

35. White, J. K. et al. Genome-wide generation and systematic phenotyping of knockout mice reveals new roles for many genes. Cell 154, 452 (2013).

36. Pettitt, S. J. et al. Agouti C57BL/6N embryonic stem cells for mouse genetic resources. Nat Methods 6, 493–495 (2009).

37. Skarnes, W. C. et al. A conditional knockout resource for the genome-wide study of mouse gene function. Nature 474, 337–344 (2011).

38. Bradley, A. et al. The mammalian gene function resource: The International Knockout Mouse Consortium. Mammalian Genome 23, 580–586 (2012).

39. Mckeown, C. R., Nowak, R. B., Gokhin, D. S. & Fowler, V. M. Tropomyosin is required for cardiac morphogenesis, myofibril assembly, and formation of adherens junctions in the developing mouse embryo. Developmental Dynamics 243, 800–817 (2014).

40. Tober, J., Yzaguirre, A. D., Piwarzyk, E. & Speck, N. A. Distinct temporal requirements for Runx1 in hematopoietic progenitors and stem cells. Development (Cambridge) 140, 3765–3776 (2013).

41. Gao, L. et al. RUNX1 and the endothelial origin of blood. Exp Hematol 68, 2–9 (2018).

42. Adamo, L. et al. Biomechanical forces promote embryonic haematopoiesis. Nature 459, 1131–1135 (2009).

43. Lundin, V. et al. YAP Regulates Hematopoietic Stem Cell Formation in Response to the Biomechanical Forces of Blood Flow. Dev Cell 52, 446–460.e5 (2020).

44. An, H. H. et al. The use of pluripotent stem cells to generate diagnostic tools for transfusion medicine. Blood 140, 10–10 (2022).

45. Goldenson, B. H., Hor, P. & Kaufman, D. S. iPSC-Derived Natural Killer Cell Therapies - Expansion and Targeting. Frontiers in Immunology vol. 13 Preprint at 10.3389/fimmu.2022.841107 (2022).

46. Thom, C. S., Chou, S. T. & French, D. L. Mechanistic and Translational Advances Using iPSC-Derived Blood Cells. J Exp Pathol 1, 36 (2020).

47. An, H. H., Poncz, M. & Chou, S. T. Induced Pluripotent Stem Cell-Derived Red Blood Cells, Megakaryocytes, and Platelets: Progress and Challenges. Curr Stem Cell Rep 4, 310–317 (2018).

48. Kolesnichenko, O. A., Whitsett, J. A., Kalin, T. V. & Kalinichenko, V. V. Therapeutic potential of endothelial progenitor cells in pulmonary diseases. American Journal of Respiratory Cell and Molecular Biology vol. 65 473–488 Preprint at 10.1165/rcmb.2021-0152TR (2021).

49. Wang, G. et al. Generation of pulmonary endothelial progenitor cells for cell-based therapy using interspecies mouse-rat chimeras. Am J Respir Crit Care Med 204, 326–338 (2021).

50. Patel, S. H. et al. Lifelong multilineage contribution by embryonic-born blood progenitors. Nature 606, 747–753 (2022).

51. Maguire, J. A., Gadue, P. & French, D. L. Highly Efficient CRISPR/Cas9-Mediated Genome Editing in Human Pluripotent Stem Cells. Curr Protoc 2, e590 (2022).

52. Kucukural, A., Yukselen, O., Ozata, D. M., Moore, M. J. & Garber, M. DEBrowser: Interactive differential expression analysis and visualization tool for count data 06 Biological Sciences 0604 Genetics 08 Information and Computing Sciences 0806 Information Systems. BMC Genomics 20, (2019).

53. Chen, E. Y., et al. Enrichr: interactive and collaborative HTML5 gene list enrichment analysis tool. http://amp.pharm.mssm.edu/Enrichr. (2013).

54. Kuleshov, M. V. et al. Enrichr: a comprehensive gene set enrichment analysis web server 2016 update. Nucleic Acids Res 44, W90–W97 (2016).

55. Xie, Z. et al. Gene Set Knowledge Discovery with Enrichr. Curr Protoc 1, e90 (2021).

56. Hu, Y. & Smyth, G. K. ELDA: Extreme limiting dilution analysis for comparing depleted and enriched populations in stem cell and other assays. J Immunol Methods 347, 70–78 (2009).

57. Korsunsky, I. et al. Fast, sensitive and accurate integration of single-cell data with Harmony. Nat Methods 16, 1289–1296 (2019).

58. Andreatta, M. & Carmona, S. J. UCell: Robust and scalable single-cell gene signature scoring. Comput Struct Biotechnol J 19, 3796–3798 (2021).

59. Schindelin, J., et al. Fiji: an open-source platform for biological-image analysis. Nat Methods 9, 676–682 (2012).

